# RNA-MobiSeq: Deep mutational scanning and mobility-based selection for RNA structure inference

**DOI:** 10.1101/2024.02.21.581340

**Authors:** Jinle Tang, Yazhou Shi, Zhe Zhang, Dailin Luo, Jian Zhan, Yaoqi Zhou

**Affiliations:** Institute of Physical and Systems Biology, Shenzhen Bay Laboratory, Shenzhen, Guangdong Province, 518107, China; Research Center of Nonlinear Science, School of Mathematical & Physical Sciences, Wuhan Textile University, Wuhan, 430200, China; School of Life and Health Sciences, The Chinese University of Hong Kong, Shenzhen, 518172, China; Ribopeutic Inc, Guangzhou International Bio Island, Guangdong, 510320, China

## Abstract

Tertiary structures of RNAs are increasingly found to be essential for their functions and yet are notoriously challenging to be determined by experimental techniques or predicted by energy-based and AlphaFold2-like deep-learning-based computational methods. Here, we coupled deep mutational scanning with mobility-based selection to separate structurally stable from nonstable RNA variants. The subsequent high-throughput sequencing allows the detection of the covariational signals of key secondary and tertiary base pairs with significantly improved, RNA-specific, unsupervised analysis of covariation-induced deviation of activity (CODA2) and amplification of these signals by Monte-Carlo simulated annealing. The resulting base-pairing structures can serve as the restraints for final energy-based structure predictions. This MobiSeq technique was tested on four structurally different RNAs. The results show that the detection of tertiary base pairs by MobiSeq allows reasonably accurate prediction of RNA tertiary structures. The proposed method should be useful for structural inference and improved mechanistic understanding of all RNAs with detectable changes in mobility due to deleterious mutations.

## Introduction

Most RNA structures remain elusive for high-resolution detection. This is because degenerated stable secondary structures and negatively charged phosphates made crystallization challenging^1^ and low number of intramolecular interactions along with six torsion angles per residue limited the application of nuclear magnetitic resonance^2^. Cryo-electron microscopy, worked well for large RNA-protein complexes, only began to address RNA-only structures^3^. As a result, only about 3% structures deposited in protein databank (https://www.rcsb.org/) contain RNAs. RNA structures has also been probed by enzymes, chemicals, hydroxyl radicals, and cross-linking, more recently, in combination with next-generation sequencing^4,5^. However, the resolution is highly variable (often not at single-nucleotide level) because of flexibility of RNA conformations, promiscuous reactivity of probes, inherent site ambiguities and sequencing noises. Moreover, unlike the success of AlphaFold in high-quality protein structure prediction by end-to-end deep learning^6,7^, the best method for RNA structure prediction techniques, according to critical assessment structure prediction techniques (CASP 15), remains traditional, energy-based global minimum searches^8,9^ by using our recently developed BRiQ statistical energy function^10^. The failure of immediate impact of deep learning in RNA structure prediction is largely due to the scarcity of RNA structural data^11^.

Recently, we showed that the full base-pairing structure of RNA, including those stabilized by tertiary interactions, can be inferred from deep mutational scanning data of self-cleavage ribozymes at the single base-pair resolution^12,13^. This was accomplished by analyzing covariation-induced deviation of activity (CODA) for both functional and non-functional variants for unsupervised classification, followed by Monte Carlo (MC) simulated annealing with mutation-derived scores. This CODA technique significantly improved the sensitivity and precision in detecting correlated mutations over these evolutionary coupling^14-16^ or epistasis^17,18^ analysis. Function-based selections, however, would be limited to the RNA with a known function that is amenable for high-throughput analysis. For a self-cleavage ribozyme, high-throughput sequencing can determine mutation-induced changes in cleavage activity by simply counting the number of sequences cleaved versus the number of sequences not cleaved^12,13^.

Here, we examined the possibility of using mutation-induced changes in mobility to separate the mutations with damaging structural changes from those without detrimental effects on RNA structures. Our hypothesis is that if a mutation denatures or partially denatures the native structure of an RNA, the mobility of the RNA mutant will be affected. As a result, structure-stable mutants can be separated from structure-destabilizing mutations by mobility-based assays (native PAGE), followed by high-throughput sequencing. Analyzing sequencing data by improved CODA (CODA2) confirmed its usefulness for inferring RNA base-pairing and three-dimensional structures.

## Results

### Overview of the RNA-MobiSeq Pipeline

The RNA-MobiSeq pipeline is a method to predict the 3D structure of a target RNA from its sequence by combining deep mutation, mobility-based separation of structurally unstable from structurally stable RNA mutants, co-evolution analysis of high-throughput sequencing data, and energy-based structure prediction. As shown in Figure 1, error-prone PCRs were first employed to generate randomly mutated sequences, which were then electro-transfected into DH5α for library constructions. The DNA libraries were further *in vitro* transcribed into RNAs that were subjected to native PAGE separation according to mobility. The bands with the same mobility as wild type RNA sequences were excised for reverse transcription. Both pre- and post-selection sequences were sent for high-throughput sequencing. The obtained sequences were then analyzed for co-variation to infer the base pairing structures using CODA2 and MC simulated annealing. The resulting base pairing structures were employed for energy minimization to model the 3D structures of the RNA by using the BRiQ statistical energy function and a folding-tree-based folding algorithm; more details can be found in Methods.

**Figure 1.**
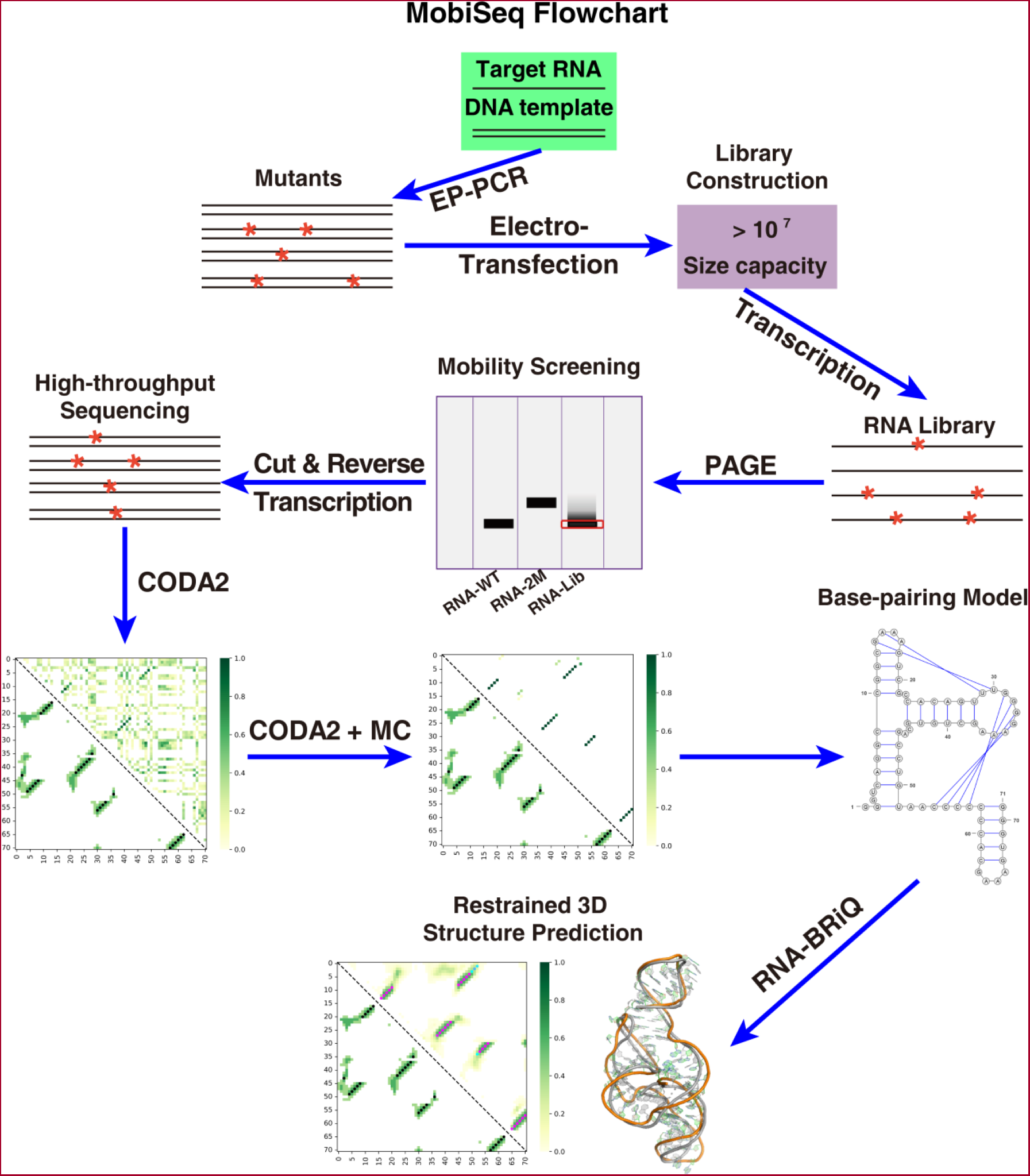
The workflow of RNA-MobiSeq for RNA 3D structure inference. The method starts from a DNA template sequence of the target RNA and employs error-prone PCR (EP-PCR) to produce a random-mutation DNA library, followed by *in vitro* transcription to an RNA library. The RNA library is, then, subjected to mobility-based selection by native polyacrylamide gel electrophoresis (PAGE). Selected RNAs are reverse transcribed for high-throughput sequencing. Sequencing results are compared to the unselected library for analyzing co-mutation signals by CODA2, following by Monte-Carlo simulated annealing for base-pairing determination. The obtained base pairs were utilized as restraints for predicting RNA structures by employing the BRiQ statistical energy function.

To test the pipeline, we chose three different types of RNAs: two riboswitches, an exonuclease resistant RNA (xrRNA), and a self-cleaving ribozyme. They are SAM-VI riboswitch from *Bifidobacterium angulatum*^19^ (PDB code: 6LAS, sequences 1-55, 55*nt*), adenine riboswitch from *Vibrio vulnificus*^20^ (PDB code: 5E54, sequences 13-83, 71*nt*, exonuclease resistant RNA from *Zika* virus^21^ (PDB code: 5TPY, sequences 1-71, 71*nt*), and inactive ribozyme mutant CPEB3-C57T^12,22^ (81 *nt*) from human (PDB code: 7QR4^23^). Their base-pairing structures and 3D structures are shown in Figures 2A, B, C, D, respectively. These RNAs were chosen for their relative small sizes and stable known three-dimensional structures. Two separate riboswitches were chosen to compare the effect of base pairing with or without tertiary-interaction-stabilized pseudoknots for adenine riboswitch and SAM-VI riboswitch, respectively. CPEB3 ribozyme was chosen so that one can compare mobility-based selection to function-based selection conducted previously^12^. All four structures are chosen very differently from each other (structural similarities between them are ∼0.1 according to TM-Score^24^, Supplementary Table S1).

**Figure 2.**
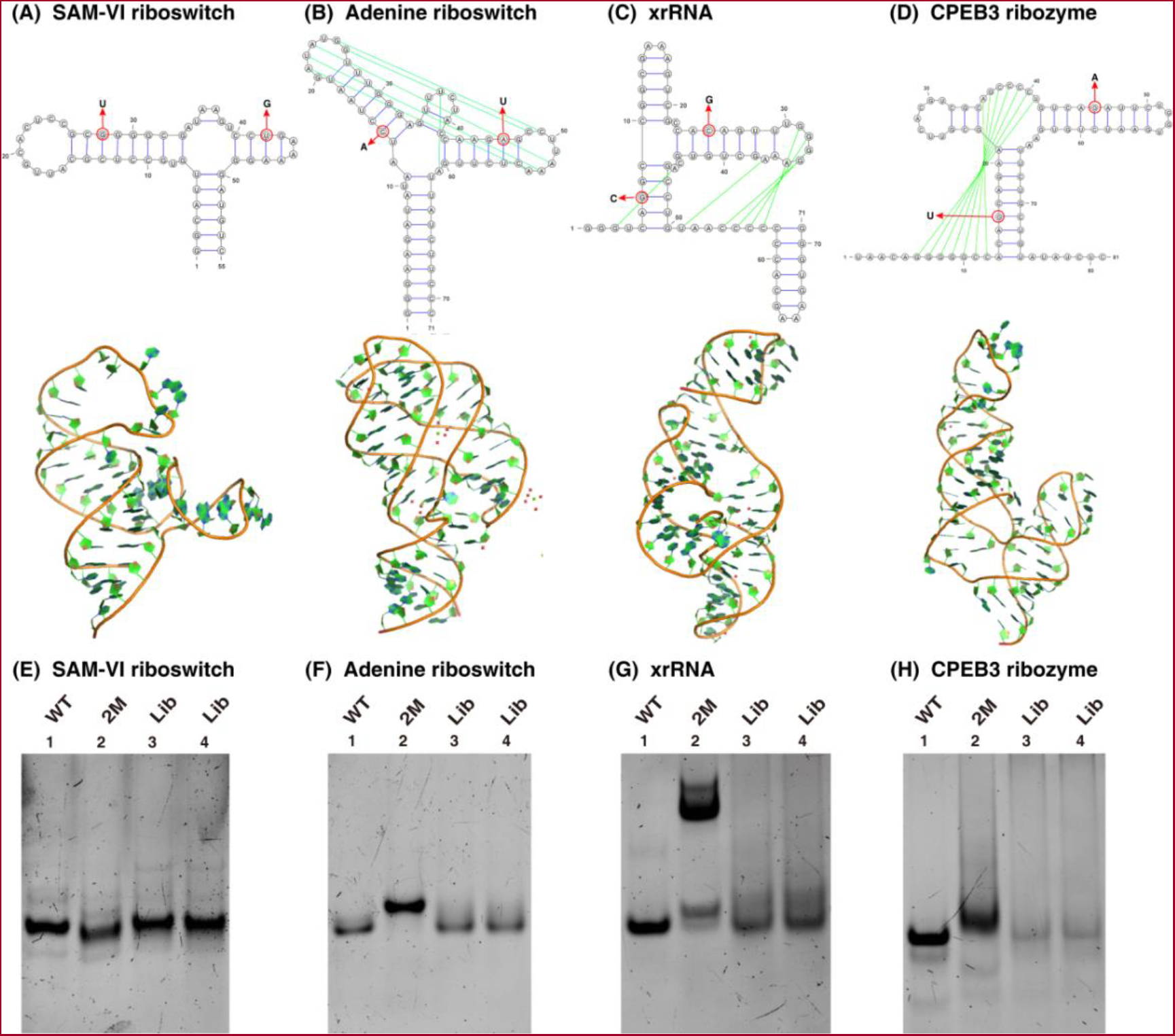
Mutation-induced Changes in Mobility as Illustrated by Four RNAs with Different Structures. The secondary and tertiary structures of SAM-VI riboswitch (PDB ID: 6LAS) (A), adenine riboswitch (PDB ID: 5E54) (B), xrRNA (PDB ID: 5TPY) (C), and CPEB3 (PDB ID: 7QR4) (D) with the locations of randomly chosen double mutants (2M) as indicated. The mobility comparison between wild type (WT), double mutants (2M), and two library replicates for SAM-VI riboswitch (E), adenine riboswitch (F), xrRNA (G), and CPEB3 ribozyme (H).

### Mobility-based Selection

To test if mutations can lead to visible change in mobility on a native PAGE, we randomly introduced two mutations for each RNA that will disrupt two base pairs in two different stems. The chosen double mutations were shown in Figure 2A for SAM-VI riboswitch, in Figure 2B for adenine riboswitch, in Figure 2C for xrRNA, and in Figure 2D for CPEB3, respectively (all RNA sequences are listed in Supplementary Table S2). Figures 2E-H compare the mobilities of wild type (WT), a disruptive double mutant (2M), and two RNA mutant library replicates generated from EP-PCR. The double mutations of all four RNAs have altered mobilities, relative to their corresponding wild-type RNAs. Meanwhile, the bands of RNA libraries are broader than the corresponding bands for the wild type. Thus, the native PAGE result indicates that the mobility is sensitive enough to separate structure-disrupting mutations from structure-unaltered mutations. Interestingly, three destabilizing RNA mutants (adenine riboswitch, xrRNA and CPEB3 ribozyme) led to slower mobility as one would expect from loosen structural forms with larger radius of gyration. However, the opposite is true for SAM-VI riboswitch. The reason for this peculiar behavior is unknown. One possible reason is that the small SAM-VI riboswitch lacks long-range tertiary interactions, leading to only small differences in the size of the disrupted structures compared to the wild type. Other effects such as charge distributions may play the role.

### Quality of Mutation Libraries

The most useful mutants for co-mutation analysis are single and double mutants. We have controlled mutation rates so that the mutation libraries are enriched with double mutations. Supplementary Table S3 shows the statistics of single, double, triple and other mutations for four RNAs before mobility-based selection. All RNA libraries have 100% coverage of single mutations and > 88% coverage of double mutations (95.2%, 88.4%, 94.4%, and 91.7% for SAM-VI riboswitch, adenine riboswitch, xrRNA, CPEB3 ribozyme, respectively). The high coverage of double mutations ensures sufficient data for our subsequent co-evolution analysis.

### Improvement of CODA2 over CODA

Both CODA and CODA2 are the methods for unsupervised detection of base pairs. There are two major differences between CODA and CODA2. First, when fitting the double-mutation fitness score using Support Vector Regression (SVR), CODA2 included mutation positions and base types before and after the mutation as the input, in addition to two single-mutation fitness values (CODA). Second, we introduced RNA-dependent SVR parameters (i.e., C andγ in SVR model), and the variance truncation coefficient *n*. These parameters were optimized for each RNA by a grid-based search for the lowest base-pairing energy (Supplementary Figure S1, see Methods for Details). As shown in Table 1 and Supplementary Figure S2, the *F1*-scores inferred from CODA2 are 0.34, 0.54, 0.64, and 0.39 for SAM-VI riboswitch, adenine riboswitch, xrRNA, CPEB3 ribozyme, respectively, which improved over CODA by 89%, 69%, 31%, and 70%, respectively. RNA-specific parameterizations are the key for significant improvement.

**Table 1.**
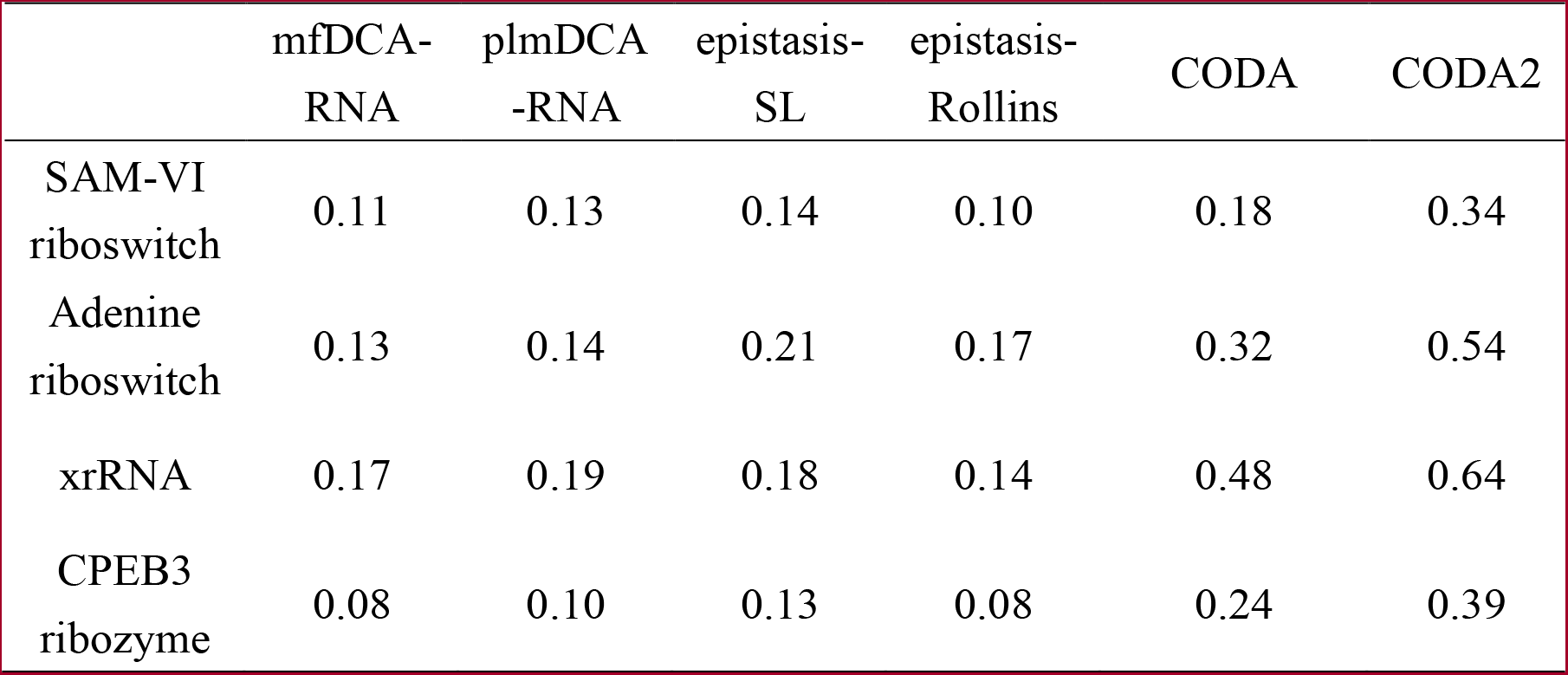
Performance of mean-field direct coupling analysis (mfDCA-RNA)^15^, pseudo-likelihood maximization coupling analysis (plmDCA-RNA)^16^, epistasis from Schmiedel & Lehner (epistasis-SL)^17^ and from Rollins *et al* (epistasis-Rollins)^18^, and covariation-induced deviation of activity (CODA in Ref. 12 and CODA2 in this work) for inferring base pairs from deep mutation data in term of *F1*-score for SAM-VI riboswitch, adenine riboswitch, xrRNA, and CPEB3 ribozyme.

|  | mfDCA-RNA | plmDCA<br>-RNA | epistasis-SL | epistasis-Rollins | CODA | CODA2 |
| --- | --- | --- | --- | --- | --- | --- |
| SAM-VI<br>riboswitch | 0.11 | 0.13 | 0.14 | 0.10 | 0.18 | 0.34 |
| Adenine<br>riboswitch | 0.13 | 0.14 | 0.21 | 0.17 | 0.32 | 0.54 |
| xrRNA | 0.17 | 0.19 | 0.18 | 0.14 | 0.48 | 0.64 |
| CPEB3<br>ribozyme | 0.08 | 0.10 | 0.13 | 0.08 | 0.24 | 0.39 |

Table 1 also presents a comparison of CODA2’s performance in inferring true base pairs from deep mutation data of four RNAs with other existing methods such as mean-field direct coupling analysis (mfDCA-RNA)^15^, pseudo-likelihood maximization coupling analysis (plmDCA-RNA)^16^, and epistasis analysis from Schmiedel & Lehner (epistasis-SL)^17^ as well as from Rollins *et al* (epistasis-Rollins)^18^, which have already been successfully applied to extract contacts from RNA families or deep mutational scanning^14-18^. Here, for method comparison, the thresholds for defining base pairs were all chosen to maximize the *F1-score*. As shown in Table 1, the four methods (i.e., mfDCA-RNA, plmDCA-RNA, epistasis-SL and epistasis-Rollins) have the similar performance in term of *F1-score* for SAM-VI riboswitch (*F1 ≤* 0.14), adenine riboswitch (*F1 ≤* 0.21), xrRNA (*F1 ≤* 0.19), CPEB3 ribozyme (*F1 ≤* 0.13), with the optimal *F1-score*s at 59%, 61%, 70%, and 67% lower than those from CODA2, respectively; see Supplementary Figure S2.

### Inference of Base-pairing Structures

Base-pairing structures can be obtained at three stages: covariation signals capture by CODA2, signal amplification by Monte-Carlo simulated annealing (CODA2+MC), and subsequent folding by BRiQ with the restraints from CODA2+MC analysis (CODA2+MC+BRiQ).

Figure 3 compares base-pairing contact maps obtained at different stages (upper triangles) with those native distance-based contact and base-pairing maps (bottom triangles) for four RNAs (SAM-VI riboswitch: Figures 3A-D, adenine riboswitch: Figures 3E-H, xrRNA: Figures 3I-L, CPEB3 ribozyme: Figures 3M-P). In these figures, CODA2 analysis yields the pairing probabilities. In addition, 1000 MC independent simulations and top 100 of 5000 independent BRiQ folding simulations were utilized to produce a distribution of pairing results because different random initial seeds will yield different outcomes. Except SAM-VI riboswitch, which does not have base pairs due to tertiary interactions, results from CODA2 can capture the signals not only from secondary base pairs but also from tertiary base pairs, including pseudoknots^25^, non-canonical base pairs and lone base pairs, which can be annotated by bpRNA^26^ or DSSR software^27^. As shown in Figures 3A-D, the structure of SAM-VI riboswitch does contain distance-based tertiary interactions. However, the lack of base pairing prohibits it having strong signals for detecting by correlated mutations. This is also true for other distance-based tertiary interactions in adenine riboswitch (Figures 3E-H), xrRNA (Figures 3I-L), and CPEB3 ribozyme (Figures 3M-P).

**Figure 3.**
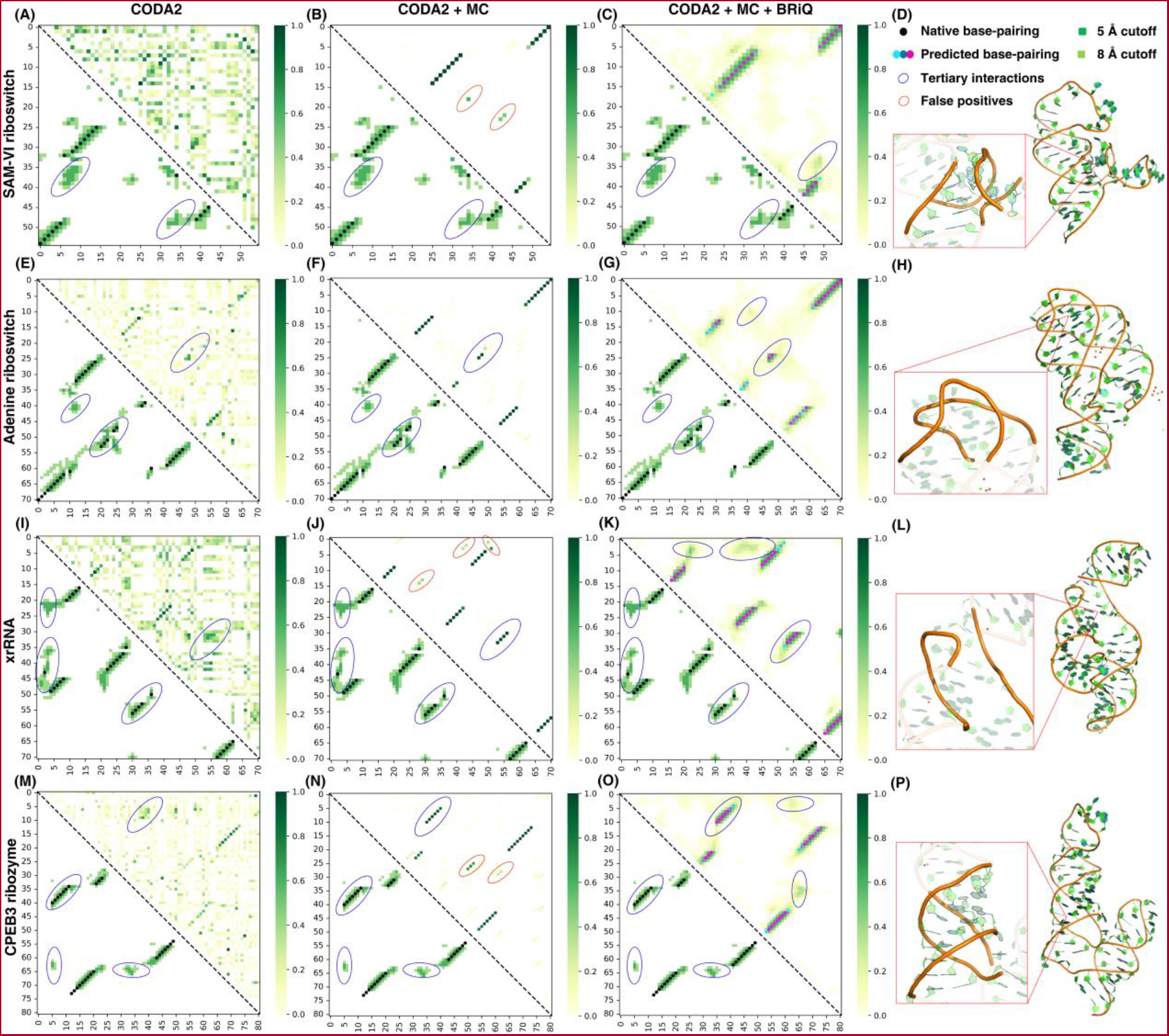
Detection of tertiary base-pairing interactions by CODA2 except for the RNA without base pairs stabilized by tertiary interactions. The base-pairing and/or distance-based contact maps (upper right) inferred from CODA2, by Monte Carlo simulated annealing (CODA2+MC), and by folding with the BRiQ statistical energy function and the restraints from CODA2+MC as well as the actual native structures (lower left) from left to right for SAM-VI riboswitch, adenine riboswitch, xrRNA, and CPEB3 ribozyme from top to bottom. Blue circles: native or predicted tertiary interactions. Red circles: false positive predictions of base-pairing. In the native/predicted distance-based contact maps, the contacts with 5 Å cutoff are shown in dark green, while those with 8 Å cutoff are shown in light green. The base-pairings from the top 100 structures predicted by BRiQ are marked by a gradient of cyan to magenta, indicating the probability from low to high. The structural domains with typical tertiary interactions in the four RNA native structures are also highlighted in the rightmost column of the figure.

Figures 3A, 3E, 3I and 3M further show that the signals from CODA2 are somewhat noisy and only those really strong signals can be detected. Monte Carlo simulated annealing builds around the base pairs with strong mutation-coupling signals in a stem to complete the stem region by an energy function (Figures 3B, 3F, 3J and 3N). Folding by the BRiQ statistical energy function further improves base pairs from CODA2+MC by removing unphysical false positives and including distance-based contacts (Figures 3C, 3G, 3K and 3O); see also in Supplementary Figure S3.

Figure 4 further compares *F1-score*s (or distribution of *F1-score*s) for four RNAs at different modeling stages to illustrate the quantitative improvement. As a reference, we also presented the results from commonly used RNA secondary structure predictor RNAfold^28^ and the folding results by BRiQ with RNAfold secondary-structure restraints. One can see that the performance of CODA2 itself may not be better than RNAfold (better for only one RNA but worse for three RNAs). This is understandable, because CODA2 can only capture those base pairs with strongest co-mutation signals as shown in Figures 3A, 3E, 3I and 3M. Adding Monte Carlo simulated annealing over the top of CODA2 quickly improves over RNAfold. For instance, the *F1-score*s for base-pairs of the best of top 5 predictions obtained from CODA2+MC are 0.89, 0.91, and 0.86 for adenine riboswitch, xrRNA, and CPEB3 ribozyme, respectively, which are 11%, 68%, and 85% higher than those from RNAfold. The only RNA, which CODA2+MC performs comparable to RNAfold, is SAM-VI riboswitch, likely due to the fact that this riboswitch does not have any base pairs stabilized by tertiary interactions and thus RNAfold can achieve unusually high performance (*F1-score* = 0.89).

**Figure 4.**
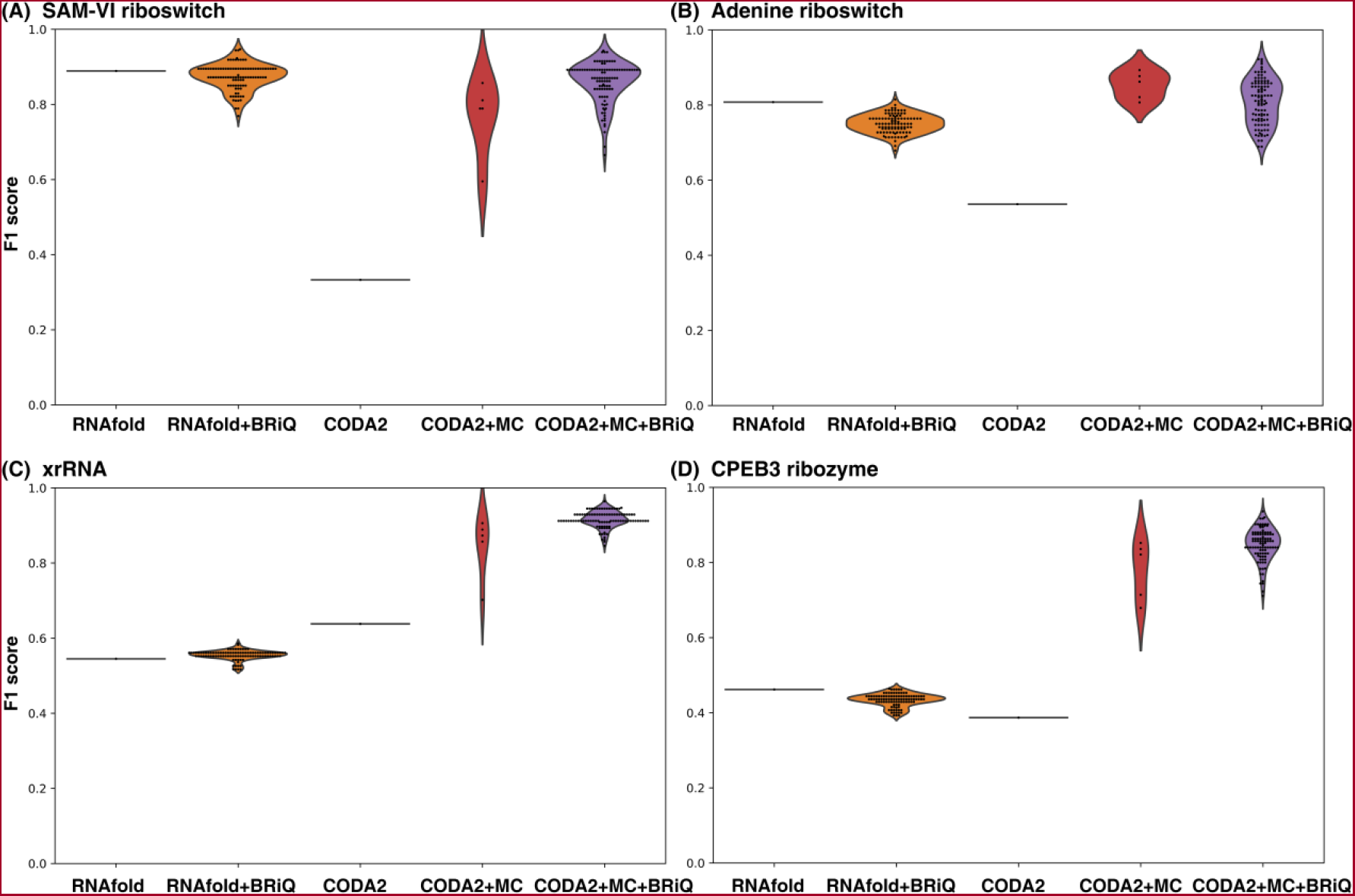
Improving base pairing results over RNAfold secondary structure prediction by Monte Carlo simulated annealing with CODA2 probabilities and folding by BRiQ with predicted base pairs as restraints. *F1-score*s for base-pairs inferred from CODA2, top five predictions from Monte Carlo simulated annealing with the CODA2 results as the seeds (CODA2+MC), and top 100 predictions from the structures folded by the BRiQ statistical energy function with restrains from the CODA2+MC base pairs (CODA2+MC+BRiQ) as compared to the reference base pairs predicted by RNAfold and top 100 predictions from the structures folded by the BRiQ statistical energy function with restrains from RNAfold predictions (RNAfold+BRiQ). (A) SAM-VI riboswitch, (B) Adenine riboswitch, (C) xrRNA, and (D) CPEB3 ribozyme.

For top 100 structures from BRiQ folding, folding with restrains from CODA2+MC yielded much better base-pairs than folding with restrains from RNAfold (mean *F1-score*s: 0.81, 0.92, and 0.85 by CODA2+MC+BRiQ versus mean *F1-score*s by RNAfold+BRiQ: 0.75, 0.55, and 0.43 with *p* value <10^−17^ for adenine riboswitch, xrRNA and CPEB ribozyme, respectively); see Figures 4B-D. The only exception is SAM-VI riboswitch (mean *F1-score*: 0.86 vs 0.87, with *p* value of ∼0.03) (Figure 4A), which does not contain base pairs stabilized by tertiary interactions. It suggests that detection of base pairs stabilized by tertiary interactions by CODA2 ensures substantially more accurate prediction of *F1-score* for adenine riboswitch, xrRNA, and CPEB3 ribozyme.

Supplementary Figure S3 further shows a comparison of base-pairs inferred from CODA2 (top 1), CODA2+MC (top 1), CODA2+MC (best in top 5), CODA2+MC+BRiQ (top1), and CODA2+MC+BRiQ (best in top 10) to native base pairs for each RNA with corresponding *F1-score*s labeled in the figure. The figure clearly demonstrates that the subsequent MC simulated annealing and BRiQ methods can effectively eliminate noise interference in the inferred base-pairing from CODA2 and complete stems with secondary or tertiary base-pairs, ultimately yielding base-pairings that closely resemble the native structure. A few tertiary base-pairs in adenine riboswitch (i.e., A_21_-A_54_, A_23_-A_52_) and xrRNA (i.e., G_3_-C_44_, A_37_-U_51_) were still not correctly inferred. The failure of CODA2 to detect non-canonical base pairs in adenine riboswitch and lone base pairs in xrRNA because their weak signals were penalized or ignored in the subsequent MC simulated annealing (see Methods).

### 3D Structures Prediction

To examine the performance on 3D structure prediction, we calculated global structural similarity scores of root-mean-squared distance (RMSD) and local distance difference test (lDDT) between predicted 3D structures and the corresponding native structure for each RNA^8,29^. We also evaluate the number of all types of steric clashes per thousand residues (clash score)^30,31^. Figure 5 compares the distributions of RMSD (0 for perfect agreement), lDDT (1 for perfect agreement), and Clash score (0, the best) for the top 100 structures folded by the BRiQ statistical energy function with restraints based on the base-pairing structures inferred from MC simulated annealing or predicted by RNAfold (Supplementary Figure S4). It is clear that 3D structures from CODA2+MC+BRiQ are substantially more accurate than those generated from RNAfold+BRiQ for the RNAs with tertiary base pairs (adenine riboswitch, xrRNA, and CPEB3 ribozymes). The improvement is even observed for SAM-VI riboswitch (*p* value < 10^−4^) despite it does not have tertiary base pairs. All predicted structures have low clash scores due to built-in penalty in the BRiQ statistical energy function.

**Figure 5.**
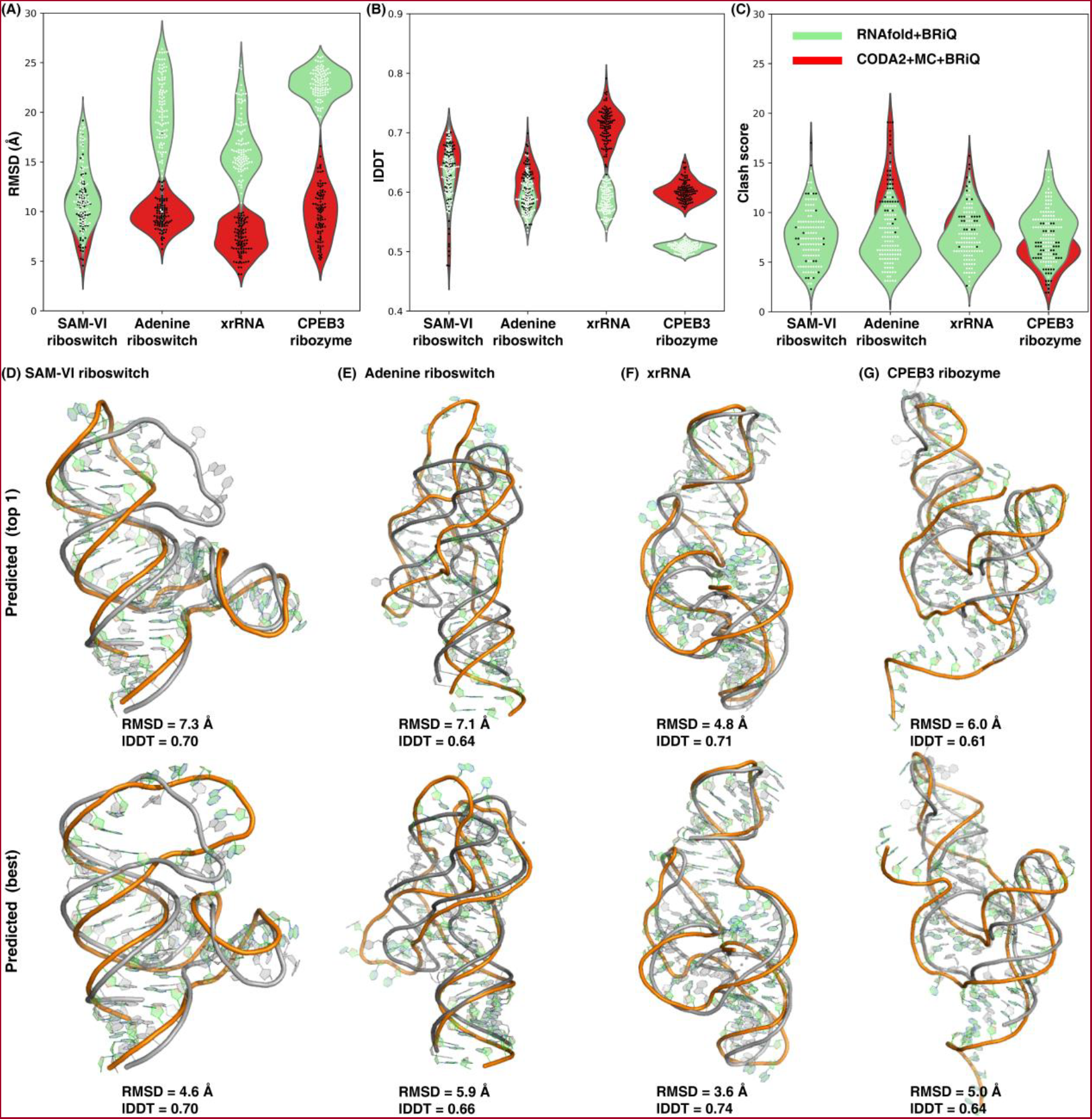
Reasonable structures obtained from BRiQ folding. The distributions of structure-prediction performance measures for four RNAs folded by BRiQ (top 100) based on the secondary structures from CODA2+MC and RNAfold, respectively, according to RMSD (A), lDDT (B), and Clash score (C). (D-G) 3D structures inferred by RNA-MobiSeq (CODA2+MC+BRiQ) for SAM-VI riboswitch (D), adenine riboswitch (E), xrRNA (F), and CPEB3 ribozymes (G). Top and bottom panels are for Top 1 and Best in Top 10, respectively. Grey: experimental structures. Color: predicted structures.

Figures 5D-G display the overlapped predicted structures with native structures. The top 1 predicted structure (according to the energy function) is 7.3 Å, 7.1 Å, 4.8 Å, and 6.0 Å in RMSD or 0.7, 0.64, 0.71 and 0.61 in lDDT scores for SAM-VI riboswitch, adenine riboswitch, xrRNA, and CPEB3 ribozyme, respectively. Best predictions in top 10 have RMSD values of 4.6 Å, 5.9 Å, 3.6 Å, and 5.0 Å in RMSD, and 0.7, 0.66, 0.74, and 0.64 in lDDT for SAM-VI riboswitch, adenine riboswitch, xrRNA, and CPEB3 ribozymes, respectively. Overall structural topology is well captured with accurate prediction for the four RNAs (Top 1 all with > 0.5 lDDT). Supplementary Figure S5 shows the change of the BRiQ energy score as a function of RMSD. All four RNAs display that a low energy score is often associated with a reasonably low RMSD value.

## Discussion

RNA structures are challenging to determine experimentally and predict computationally. Here, we propose a simple and yet effective method called MobiSeq that integrates an age-old molecular biology technique with the modern computational method for RNA structure inference. More specifically, we combine deep mutational scanning and mobility selection with high throughput sequencing. The resulting sequencing data is then subjected to covariation analysis by the newly developed method CODA2, Monte Carlo simulations, and base-pair-restrained structure prediction by the BRiQ energy function.

We have previously performed deep mutational scanning of CPEB3 ribozyme based on its self-cleavage activity^12^. It is of interest to compare function-based and mobility-based selection results for CPEB3. The double mutation coverage was only 41% by function selection, compared to 92% in this work. Despite the poorer coverage, function-based selection leads to much higher performance of base pair inference (0.55 versus 0.24 for *F1-score* by CODA). The *F1-score* of CPEB3 ribozyme for mobility-based screening reaches 0.39 only after using a more sensitive CODA2, which is still far below 0.55. More importantly, previous CODA analysis can capture tertiary lone base pairs and non-canonical base pairs of CPEB3 ribozyme and twister ribozyme, but none for CODA2 analysis for three RNAs with those tertiary interactions (Supplementary Figure S3B-D). This poorer performance for MobiSeq is largely because mobility-based selection requires manual cut of the gel based on the location of wild types. As a result, it lacks the precision for the selection based on self-cleavage activity, which can be directly measured by counting the number of cleaved and non-cleaved sequences from high-throughput sequencing data. Although the quality of mobility-selected mutation library is not as high as that of the function-selected library, the key pseudoknot in CPEB3 is captured by the mobility selection methods as shown in Supplementary Figure S3D.

Moreover, MobiSeq did not just capture pseudoknots in CPEB3 ribozyme. It identifies a pseudoknot for all three RNAs with pseudoknots studied in this work (Supplementary Figure S3B-D). Thus, MobiSeq offers reliable new technique for detection of tertiary interactions of all RNAs whose mutations can affect their mobilities. The detection of the signal for tertiary base pairs by CODA2 allows subsequent improvement of base pair prediction by Monte Carlo simulated annealing. The resulting restraints lead to far more accurate 3D structures than the baselines from BRiQ-based folding with RNAfold secondary-structure restraints. Thus, MobiSeq offers a new general method, alternative to experimental structure determination, to detecting tertiary interactions.

Certainly, the performance of MobiSeq for 3D RNA structure prediction has not yet reached the high-accuracy level achieved by AlphaFold2 for proteins. In the recent CASP 15, the CPEB3 ribozyme (i.e., target R1107) was used to assess RNA modeling. The top 5 models are from AIchemy_RNA2 group, with lDDT of 0.72, 0.72, 0.71, 0.73, and 0.72, which are about 0.1 higher than the top 5 model from MobiSeq (0.61, 0.60, 0.65, 0.61, and 0.62). This improvement could be attributed to their effective utilization of the known homologous template (i.e., PDB: 4PR6, with sequence identity of 62.3%)^9^. However, the average lDDT (0.62) of the top 5 models inferred by MobiSeq is significantly higher than the overall mean lDDT (0.44) of all 131 models in CASP 15 (see https://predictioncenter.org/casp15/results.cgi?tr_type=rna). In addition to CPEB3 ribozyme, the average lDDT values of the top 5 models predicted by CODA2+MC+BRiQ for SAM-VI riboswitch, adenine riboswitch, and xrRNA are 0.65, 0.65, and 0.72, respectively. That is, the mean lDDT of top 5 models for the four RNAs is 0.66, which is ∼40% higher than the mean lDDT (0.47) of 153 models submitted for the smaller PreQ1 class I type III riboswitch (i.e., target R1117, H-type pseudoknot with 30nt length) in CASP15. The results suggest that MobiSeq has the ability to accurately infer RNA tertiary structures based on the deep mutational scanning and mobility-based selection, without relying on structure templates or manual intervention.

The reasonable but not yet high-accuracy prediction of RNA structures is largely due to limitation of the BRiQ energy function. For a perfect energy function, a native structure should have the lowest energy. We utilized BRiQ to optimize the native structures of SAM-VI riboswitch and xrRNA and compared them with the predicted 3D structures from CODA2+MC+BRiQ (We did not perform this optimization for adenine riboswitch and CPEB3 ribozyme because the sequences in PDB structures are slightly different.). As shown in Supplementary Figures S5A and S5C, the BRiQ refinement did not result in significant changes to the native structure for SAM-VI riboswitch and xrRNA (RMSD < 1.5 Å). However, the energies of refined native structures are not significantly lower than the energies of the top structures predicted by CODA2+MC+BRiQ. Thus, the BRiQ potential does not yet achieve global minima for all native structures. In other words, the failure to locate native structures is not due to trapping in local minima but due to the failure of the energy function to make native structures as global minima for most cases. For proteins, AlphaFold2 successfully overcomes this energy-function problem by performing end-to-end training^7^. Such AI-based advance is yet to achieve for RNA structure prediction due to lack of sufficient RNA structures^11^.

It is worth mentioning that it takes about one month for EP-PCR library construction, mobility-based screening, and HTS sample preparation after the genes synthesized. It often took 3 rounds of EP-PCR to achieve high coverage of double mutations. The turn-around-time frame can be further cut to 2-3 weeks if doped libraries, rather than EP-PCR were directly employed to produce mutation libraries. The gene synthesis, materials, consumables, and sequencing costs take < $300 (US dollars) per RNA. Thus, MobiSeq is not only an effective but also inexpensive technique for tertiary interaction detection.

## Materials and Methods

### DNA Templates, RNA Synthesis, and Purification

The corresponding DNA templates of four RNAs were directly synthesized and subcloned into pUC57-1.8k vector with N-terminal T7 RNA polymerase promoter/ adaptor sequence-1 and C-terminal adaptor sequence-2/BspQ I restriction enzyme site (General Biosystems, China). The detailed sequence information was listed in Supplementary Table S2. All plasmids were amplified in DH5α and extracted using Plasmid Mini Kit I (Omega, D6943-02).

The restriction enzyme BspQ I was utilized to linearize the plasmid. Linearized vectors were recycled with Gel Extraction Kit (Omega, D2500-02) and eluted with nuclease-free water. Then, linearized vectors were employed as templates for *in vitro* RNA synthesis by T7 RNA polymerase at 37 °C for 5 h. The reaction system was as follows: 20 μL volume containing 1× reaction buffer (NEB, B9012S), 5 U/μL T7 RNA polymerase (NEB, M0251L), 1 U/μL RNase inhibitor (NEB, M0314S), 1 mM NTPs (NEB, N0416S) and DNA template. After transcription, DNase I (NEB, M0303L) was added to the reaction system for further incubation at 37 °C for 30 mins to eliminate the DNA template. The synthesized RNAs were purified according to the manufacturers’ instructions of RNA Clean & ConcentratorTM-5 (Zymo Research, R1016) and finally eluted with nuclease-free water. The concentrations of RNAs were determined using NanoDrop™ One (Thermo Scientific). The purity of all newly synthesized RNAs was detected using UREA-polyacrylamide gel electrophoresis (PAGE) which was stained with SYBR™ Gold (Invitrogen, S11494). The gels were finally visualized on a ChemiDoc MP imaging system (Bio-Rad).

### Preparation of DNA Mutation Libraries

Error-prone PCR (EP-PCR) is a method commonly employed to introduce random mutations into a defined segment of DNA and generated gene variant libraries for directed evolution^32^. The wildtype sequences of selected RNAs were randomly mutated using a QuickMutation™ Random Mutagenesis Kit (Beyotime, D0219S) with primers M13-FP and M13-RP (Supplementary Table S4). A total of three rounds of EP-PCR were performed for random mutation of the original sequence. Every round PCR products were separated on 10% UREA-PAGE and the target bands with expected sizes were excised, crushed and frozen at -80 °C. The frozen solidified products were quickly thawed in a 90 °C hot bath for 5 mins and the target DNA was passive eluted in TE/NaCl buffer (10 mM Tris–HCl, 1mM EDTA, pH 8.0, 100 mM NaCl) on a rotary shaker at 37 °C for overnight. The DNAs were finally precipitated using ethanol and redissolved in nuclease-free water. The recycled target DNA products were cloned into pUC57-1.8k vector using ClonExpress II One Step Cloning Kit (Vazyme, C112-02) and transformed to DH5α for Sanger sequencing for calculating the mutated rate.

Based on the obtained mutated rate,the third round PCR products were chosen for DNA library construction. The fragments of the mutated DNA library were amplified with primers M13-Plus-FP/M13-Plus-RP and the linearized pUC57-T7Q vector was amplified with primers AS-FP/AS-RP using PrimerSTAR HS DNA polymerase (Takara, R040A) (Supplementary Table S4). The DNA fragments were loaded on the agarose-gel and the target bands were excised and recycled using a Gel Extraction Kit (Omega, D2500-02). DNA fragments and pUC57-T7Q vector were mixed with Exnase II in the reaction buffer on ice and the reaction system was incubated at 37 ? for 30 min (CloneExpress II One Step Cloning Kit, Vazyme, C112-02). The assembled products were purified with the RNA Clean & Concentrator™-5 (Zymo Research, R1016) and eluted with nuclease-free water.

The electrocompetent DH5α was prepared as follows. One DH5α colony was picked and cultured in 5 mL LB medium at 37 ? for overnight. One milliliter of overnight cultured DH5α was pipetted and transferred into 200 mL LB medium and continued to be cultured at 37 °C. Cells were harvested using centrifugation when the density of OD600 reached 0.6. The DH5α pellet was washed one time using ice-cold ultrapure-water and three times using ice-cold 10% glycerol buffer. Finally, electrocompetent cells DH5α was resuspended in 2 mL 10% glycerol buffer for further electro-transformation. The assembled products were mixed with electrocompetent cells and transformed into electrocompetent cells in a 2-mm Gene Pulser cuvette (Bio-Rad, 165-2086) using an electroporation instrument (Bio-Rad, MicroPulser). The transformants were cultured on 20 mL LB medium containing 100 μg/mL ampicillin at 37 °C for 10 h and collected using centrifugation. The pellet was resuspended in LB medium containing 15% glycerol and stored at -80 °C for long term use. The size of the constructed library was detected by the number of colonies after a gradient dilution on LB-agar-Amp plate. Ten clones were randomly selected and sent for Sanger sequencing to evaluate the percentage of clones with the right insertion form in our libraries. The size capacities of constructed libraries were all greater than 1 × 10^7^.

### Mobility Screening of RNA Sequences by Native-PAGE

Native PAGE is a well-established method for RNA conformation and folding probing^33^. In native gels, RNA mobility is determined by conformation, folding and size of the RNA^34^. As the size of RNA sequences in library is the same, so the mutated RNA sequences with the same mobility as the wild type RNA sequence could be screened by native PAGE. The sizes of 10^7^ clones were taken out from the mutated library, respectively, cultured, and amplified. The corresponding plasmids were extracted. The BspQ I linearized vectors were transcribed to RNAs. RNA samples were heated to 70 °C for 5 min and slowly cooled to 4 °C in the folding buffer containing 10 mM MgCl2, 10 mM KCl, 10 mM sodium acetate, and pH 6.8. RNA samples were mixed with nondenaturing loading buffer and were finally loaded to native page gels for running at 60 V for 10 - 12 h at ice bath equipment in a cold room. The bands which showed the same mobility as wild type RNA sequences were excised and RNAs was extracted and purified used the method as DNA extraction in PAGE gel, as described above.

### Sequencing Library Preparation and High-throughput Sequencing (HTS)

The purified RNA was reverse transcribed using ProtoScript® II Reverse Transcriptase (NEB, M0368L). The 20 μL reaction volume containing 4 µL 5 × ProtoScript II buffer, 2 μL 0.1 M DTT, 1 μL 10 mM dNTP, 0.2 μL RNase inhibitor (40 U/μL), 1 μL ProtoScript II RT (200 U/µL), 2 μL M13F-Plus-RP and the RNA template. The reaction system was incubated at 42 °C for 1 h and then inactivated at 65 °C for 20 min. The cDNA product was stored at -20 °C for further use. 10 μL cDNA product was utilized as the template for amplification (10 cycles) of the fragments with primers M13-FP/M13-RP. Then, the DNA sequence library was constructed using recycled fragments as the template with adaptor primers (Supplementary Table S4) in PCR amplification reactions. The PCR products were purified by agarose gel electrophoresis with a Gel Extraction Kit (Omega, D2500-02). The DNA concentration was measured using Qubit (Invitrogen™) and peak size was analyzed by Agilent 2100-H. The DNA samples were high throughput sequenced using Illumina NovaSeq at Novogene.

### HTS Data Analysis

High-throughput sequencing data were obtained from Novogene Technology Co., Ltd using NovaSeq PE150. Sequencing data analysis was similar to our previous work on self-cleaving ribozymes^12^. Paired-end sequencing reads were first merged into single reads by using PEAR^35^. Then, we counted the numbers of reads of all the mutants and the wild type pre- and post-native PAGE separation. Because an RNA mutant will change the balance between the frequency at the native structure and the frequency at other structures, we can calculate a mutant *i*’s fitness *fi* according to their read counts at pre- and post-native PAGE separation (*n*_*i*_^*pre*^ and *n*_*i*_^*post*^, respectively), relative to wild-types (*n*_*wt*_^*pre*^ and *n*_*wt*_^*post*^, respectively), i.e.,

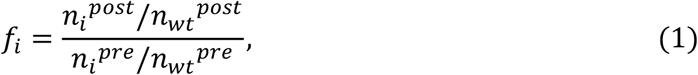

### Improved CODA Analysis for Mobility-selected RNAs

CODA analysis assumes that the effects of two mutations on an RNA structure are independent of each other if they are not in close contact^12^. In this case, the fitness of the double mutant of these two mutations can be predicted by the fitness of two single-mutants. On the other hand, for two pairing bases, the fitness of their double mutations will be higher than what could be predicted from two independent single mutations because single mutations will disrupt and double mutations will restore the structure. Thus, the base-base interaction could be detected by examining covariation-induced deviation of the fitness score of a double-mutant from the independent mutation model^12^.

To establish an independent-mutation model, we built a dataset for all double mutants including eight features for each double mutant (the sequence positions of two mutations (*pos*), wild (*wt*) and mutational (*mt*) base types, and two fitness scores of two single mutants) and one label (the fitness score of the double mutant). Base types were encoded with one-hot encoding (1000, 0100, 0010, and 0001 for A, C, G, and U, respectively). After max-min normalization of the data, the model was constructed using a support vector machine regression (SVR) to fit the fitness scores of all double-mutation variants as a function of the features of their corresponding single-mutation variants, that is, *f*(*X Y*) = (*pos*(*X*) *pos*(*Y*) *wt*(*X*) *wt*(*Y*) *t*(*X*) *t*(*Y*) *f*(*X*) *f*(*Y*)), where *X* and *Y* are the two corresponding single-mutants in the double mutant. Then, the fitness score for any double mutants can be predicted by the trained SVR model, and the CODA score was calculated through the deviations between predicted (*f*_*pred*_) and actual (*f*_*act*_) fitness of the double-mutant, CODA = (*f*_*act*_ − *f*_*pred*_)/(*f*_*pred*_ + 0.2), where the shift of 0.2 is employed to avoid near zero value of *f*_*pred*_^12^. This regression equation is different from the original CODA by including mutation positions and base types before and after mutations as the features.

We expected that the distribution of CODA scores for double mutations is a mixture of two distributions approximated as Gaussian. Due to the dominance of unpaired bases, the first Gaussian distribution of CODA score was fitted from all double mutation data. Then, the second distribution was obtained by the subset of data points with CODA score > *μ*_1_ + *nσ*_1_, based on the expectation (*μ*_1_) and standard deviation (*σ*_1_) of the first distribution, where *n* is an adjustable parameter; see Supplementary Figure S6. Finally, the pairing score (*PS*) of a double mutation was calculated by a two-state naive Bayesian classifier with

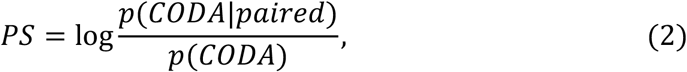

where *p*(*CODA*) = *p*(*paird*) × *p*(*CODA*|*paired*) + (1 − *p*(*paird*)) × *p*(*CODA*| *unpaired*), and the probability is derived from the mixture Gaussian model, without using a training set^12^.

Since the parameters (i.e., C and γ) in SVR, as well as the above adjustable parameter *n* (i.e., the variance truncation coefficient) generally depend on the distribution of different RNA mutation data, we performed a grid search on the parameters for each data set, and calculated the energy of inferred base-pairing under different parameter settings using the thermodynamic parameters36 (see Eq. 3 in the next section). The contacts with the lowest free energy are then selected as the final inference result; see Supplementary Figure S1.

### Base-pairing Prediction by Monte Carlo Simulated Annealing (CODA2+MC)

As in our previous work^12^, we further enhanced pairing signals by integrating the above pairing score *PS* with experimental free energies for stacking by

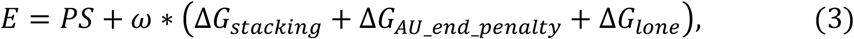

where ω is an optimized weight (0.9 in this work). Δ*G*_*stacking*_ in Eq. (3) is the sequence-dependent stacking energy between two adjacent base pairs, calculated by Δ*G*_*stacking*_= Δ*H*_*stacking*_ − *T*Δ*S*_*stacking*_, where Δ*H*_*stacking*_ and Δ*S*_*stacking*_ are from the thermodynamic measurements^35^, and *T* is real temperature in Kelvin.

Δ*G*_*AU end penalty*_ is the penalty of each helical AU end^36^, and Δ*G* _*lone*_ (0.5kcal/mol) is the penalty for a lone base pair. Unlike the previous work^12^, we did not add a penalty for noncanonical base pairs but instead ignore them in MC simulated annealing for a smoother energy landscape.

In MC simulated annealing, the initial temperature was set at 120? and decreased by a factor of 0.95 until temperature reached ∼25?. At each temperature, 1×10^5^ MC steps were performed. In each MC step, a random new base pair (*i-j*) between bases *i* and *j* (|*j-i*|>3) was generated while the existing pairs related to the new pair were destroyed, and the change is accepted or rejected according to the Metropolis criterion^37^. At the end of the simulated annealing procedure, the secondary structure with the lowest energy was regarded as the prediction. This annealing procedure could run independently multiple times (e.g., 1000), and after clustering, the top 5 base-pairing structures were taken as the final predicted results.

### Tertiary Structure Prediction Using the RNA-BRiQ Statistical Potential

We further employed the RNA-BRiQ statistical energy function^10^ for building 3D structures for RNAs. The base-pairing structures predicted above (CODA2+MC) from deep mutational scanning data as well as the sequence of an RNA were inputted for energy minimization by a nucleobase-centric tree algorithm with default parameters. For each of top 5 predicted base-pairings of an RNA, 1000 3D models were constructed using the RNA-BRiQ with different random seeds. Then, all the 3D models for the top 5 base-pairing structures (i.e., 1000×5) were further scored by the BRiQ energy, and the 100 models with the lowest energy scores (i.e., top 100) were taken as the final predicted tertiary structures.

### Performance Evaluation

The predicted contact maps (from CODA2) as well as the base-pairing (from MC or BRiQ) were evaluated by *F1-score* (*F*1), which is the harmonic mean of sensitivity (*Sen*) and precision (*Pre*) and defined by

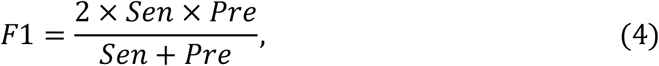

where *Sen* =*TP*/(*TP* + *FN*) and *pre* = (*TP* + *FP*). Here, *TP, FN, FP* and *TN* are the numbers of true positive, false negative, false positive and true negative contacts in the prediction, respectively. The base pairs of native structures and structures predicted by BRiQ were derived by the DSSR software^27^, and the maximal *F1* was used to characterize the accuracy of the top 1 predicted contacts from CODA2 with different parameters.

As in CASP 15, we evaluated 3D predicted structures by RMSD (root mean square deviation) and lDDT (local distance difference test)^29,38^. The lDDT (unlike for proteins, the implementation of lDDT for RNA did not penalize for stereochemical violations) was calculated by OpenStructure software (http://www.openstructure.org)^38^. In addition, we employed RNA puzzles metrics such as clash score for assessing the atomic harmony of the structure^31^. The clash score was calculated by MolProbity (http://molprobity.biochem.duke.edu/)^30^.

### Mutational coupling analysis and epistasis

Functional variants with high fitness score were used as the input sequences for covariation analysis by mfDCA-RNA and plmDCA-RNA; see more details in Ref. 12. The pydca, a standalone Python-based software package for the DCA of protein- and RNA-homologous families (https://github.com/KIT-MBS/pydca)^39^ was used to infer coevolutionary couplings between pairs of nucleotides, based on two popular inverse statistical approaches, namely, the mean-field (mfDCA-RNA) and the pseudo-likelihood maximization (plmDCA-RNA) with default parameters.

The epistasis was calculated from a non-parametric null model following Schmiedel & Lehner (epistasis-SL)^17^ and Rollins et al. (epistasis-Rollins)^18^. In epistasis-SL, the combined score from the three interaction scores was used to estimate the epistatic interactions for the four RNAs (https://github.com/jschmiedel/DMS2structure).

## Data Availability

Illumina sequencing data generated in this study are deposited in the NCBI Sequence Read Archive (SRA) under BioProject accession number PRJNA1001602. The codes as well as scripts needed to repeat the analyses are available at GitHub: https://github.com/RNA-folding-lab/CODA2.

## Acknowledgement

This research received support from National Key Research and Development Program of China (NO.2021YFF1200400) and the National Natural Science Foundation of China (Grant #22350710182) and benefited from access to the supercomputing resources at Shenzhen Bay Laboratory.

## Conflict of Interest

All authors declare no financial interest. Zhan is the founder and CEO and Zhou is the scientific founder for Ribopeutic, respectively.

## Author Contributions

JZ and JT conducted experimental design. JT performed wet-lab experiments. YS performed data analysis, computational prediction and method development. ZZ and DL participated initial data analysis and method development. JZ and YZ initiated and supervised the project. YZ provided the funding support. JT, YS, and YZ drafted the whole manuscript. All authors involved subsequent manuscript improvement.

## Notes

### Competing Interest Statement

The authors have declared no competing interest.

