## Supplementary materials for "RNA-MobiSeq: Deep mutational scanning and mobility-based selection for RNA structure inference"

**Supplementary Table S1**. The structural similarity according to TM-score between four RNAs employed in this work.

| RNAs | SAM-VI riboswitch | Adenine riboswitch | xrRNA | CPEB3 ribozyme |
|---|---|---|---|---|
| SAM-VI riboswitch | 1.0 | 0.102 | 0.036 | 0.106 |
| Adenine riboswitch | 0.086 | 1.0 | 0.044 | - [a] |
| xrRNA | 0.028 | 0.042 | 1.0 | 0.042 |
| CPEB3 ribozyme | 0.094 | - | 0.044 | 1.0 |

[a] There is no alignment between two RNAs.

**Supplementary Table S2.** The DNA template sequences of native RNAs as well as the designed double mutations (denoted as 2M).

| Name [a] | | Sequence (5'-3') [b] |
|---|---|---|
| SAM-VI riboswitch | WT | GGCATTGTGCCTCGCATTGCACTCCGCGGGGCGATAAGTCCTGAAAAGGGATGTC |
| | 2M | GGCATTGTGCCTCGCATTGCACTCCGCTGGGCGATAAGTCCGGAAAAGGGATGTC |
| Adenine riboswitch | WT | GGGAAGATATAATCCTAATGATATGGTTTGGGAGTTTCTACCAAGAGCCTTAAACTCTTGATTATCTTCCC |
| | 2M | GGGAAGATATAATACTAATGATATGGTTTGGGAGTTTCTACCTAGAGCCTTAAACTCTTGATTATCTTCCC |
| xrRNA | WT | GGGTCAGGCCGGCGAAAGTCGCCACAGTTTGGGGAAAGCTGTGCAGCCTGTAACCCCCCCACGAAAGTGGG |
| | 2M | GGGTCACGCCGGCGAAAGTCGCCAGAGTTTGGGGAAAGCTGTGCAGCCTGTAACCCCCCCACGAAAGTGGG |
| CPEB3 ribozyme | WT ($C_{57}T$) | TAACAGGGGGCCACAGCAGAAGCGTTCACGTCGCAGCCCCTGTCAGATTCTGGTGAATCTGTGAATTCTGCTGTATATCTC |
| | 2M | TAACAGGGGGCCACATCAGAAGCGTTCACGTCGCAGCCCCTGTCAAATTCTGGTGAATCTGTGAATTCTGCTGTATATCTC |

[a] "-WT" represents the DNA template for the native RNA sequence; "-2M" represents the DNA template for an disruptive double mutant shown in Figure 2 in the main text. [b] The mutant bases are marked with red color. For CPEB3 ribozyme, in order to facilitate mobility screening, the 57-th base (i.e., C) was also mutated into T (marked with green color) to reduce their activity.

**Supplementary Table S3**. The number of variants and the coverage of single, double, triple and other mutations of the four RNAs examined.

| RNAs | Length (nt) | Mutations [a] | Number of variants | Mutation coverage [b] |
|---|---|---|---|---|
| SAM-VI riboswitch | 55 | 1 | 165 | 100% |
| | | 2 | 12722 | 95.2% |
| | | 3 | 24966 | 3.5% |
| | | 4-9 | 69 | - |
| Adenine riboswitch | 71 | 1 | 213 | 100% |
| | | 2 | 19770 | 88.4% |
| | | 3 | 4546 | 0.29% |
| | | 4-8 | 86 | - |
| xrRNA | 71 | 1 | 213 | 100% |
| | | 2 | 21121 | 94.4% |
| | | 3 | 89575 | 5.8% |
| | | 4-10 | 38020 | - |
| CPEB3 ribozyme | 81 | 1 | 243 | 100% |
| | | 2 | 26741 | 91.7% |
| | | 3 | 66277 | 2.9% |
| | | 4-10 | 28032 | - |

[a] The number of mutated bases in each sequence. Because the sequences with multiple mutations are relatively fewer (as we were more interested in double mutations), we have consolidated multiple mutations (>3) together. [b] "-" indicates that the mutation coverage is very low (~0%).

**Supplementary Table S4**. The primers used in this study.

| Name | Sequence (5’ - 3’) | Notes |
| --- | --- | --- |
| M13-FP | GGTTTTCCCAGTCACGAC | EP-PCR primer |
| M13-RP | GTAAAACGACGGCCAGTG | EP-PCR primer |
| M13-Plus-FP | TAGCGGTTTTCCCAGTCACGAC | Amplification of EP-PCR products for library construction |
| M13-Plus-RP | CTTCTGTAAAACGACGGCCAGTG | Amplification of EP-PCR products for library construction |
| AS-FP | CACTGGCCGTCGTTTTACAGAAGAGCacat | Amplification of vector for library construction |
| AS-RP | GTCGTGACTGGGAAAACcgctatagtgagtcgtattag | Amplification of vector for library construction |
| P5-089-FP | AATGATACGGCGACCACCGAGATCTACACACACTAAGACACTCTTTCCCTACACGACGCTCTTCCGATCTgGTTTTCCCAGTCACGAC | For NGS Samples preparation |
| P5-090-FP | AATGATACGGCGACCACCGAGATCTACACGTGTCGGAACACTCTTTCCCTACACGACGCTCTTCCGATCTgGTTTTCCCAGTCACGAC | For NGS Samples preparation |
| P5-091-FP | AATGATACGGCGACCACCGAGATCTACACTTCCTGTTACACTCTTTCCCTACACGACGCTCTTCCGATCTgGTTTTCCCAGTCACGAC | For NGS Samples preparation |
| P5-092-FP | AATGATACGGCGACCACCGAGATCTACACCCTTCACCACACTCTTTCCCTACACGACGCTCTTCCGATCTgGTTTTCCCAGTCACGAC | For NGS Samples preparation |
| P7-089-RP | CAAGCAGAAGACGGCATACGAGATGTGCGATAGTGACTGGAGTTCAGACGTGTGCTCTTCCGATCTGTAAAACGACGGCCAGTG | For NGS Samples preparation |
| P7-090-RP | CAAGCAGAAGACGGCATACGAGATACATAGCGGTGACTGGAGTTCAGACGTGTGCTCTTCCGATCTGTAAAACGACGGCCAGTG | For NGS Samples preparation |
| P7-091-RP | CAAGCAGAAGACGGCATACGAGATGAACATACGTGACTGGAGTTCAGACGTGTGCTCTTCCGATCTGTAAAACGACGGCCAGTG | For NGS Samples preparation |
| P7-092-RP | CAAGCAGAAGACGGCATACGAGATAGGTGCGTGTGACTGGAGTTCAGACGTGTGCTCTTCCGATCTGTAAAACGACGGCCAGTG | For NGS Samples preparation |

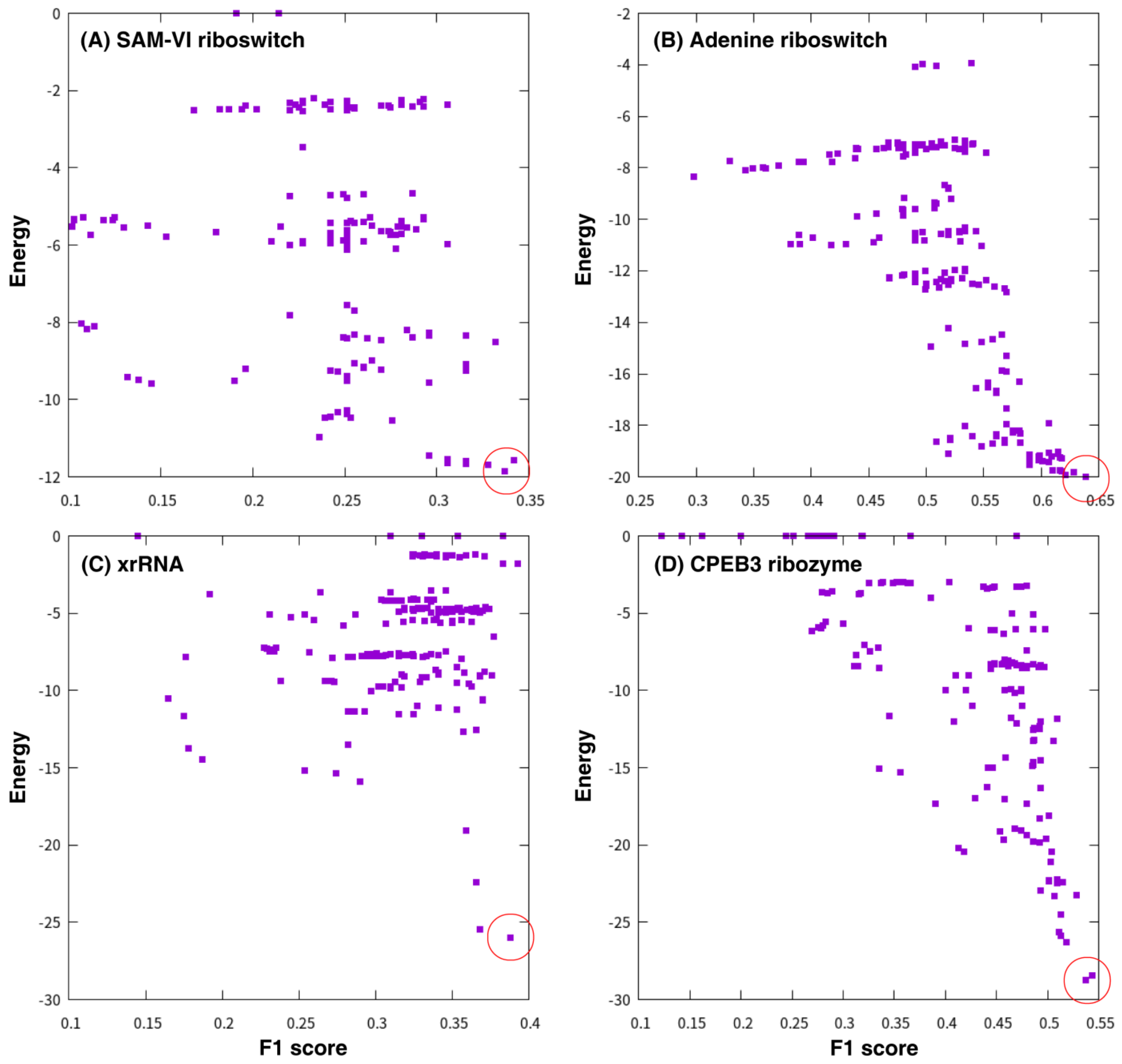


**Supplementary Figure S1. RNA-specific CODA2 parameters.** Unlike the original version of CODA[1], to increase the adaptability of the model to different data distributions, we iterated through several adjustable parameters (i.e., C and $\gamma$ in SVR model, and the variance truncation coefficient $n$) to obtain a series of contact maps. These maps were further scored by the energy calculated by Eq. 3 in the main text using the experimental thermodynamic parameters[2], and the one with the lowest energy was taken as the prediction. (A-D) Energy versus *F1* score of the maps inferred by CODA2 with different parameters for the four RNAs: SAM-VI riboswitch (A), Adenine riboswitch (B), xrRNA (C), and CPEB3 ribozyme (D). The $(C, \gamma, n)$'s for the results with lowest energy are (1.0, 1.0, 2.5), (0.01, 0.1, 3.5), (5.0, 10.0, 3.0), and (100.0, 1.0, 4.0) for SAM-VI riboswitch, Adenine riboswitch, xrRNA, and CPEB3 ribozyme, respectively.

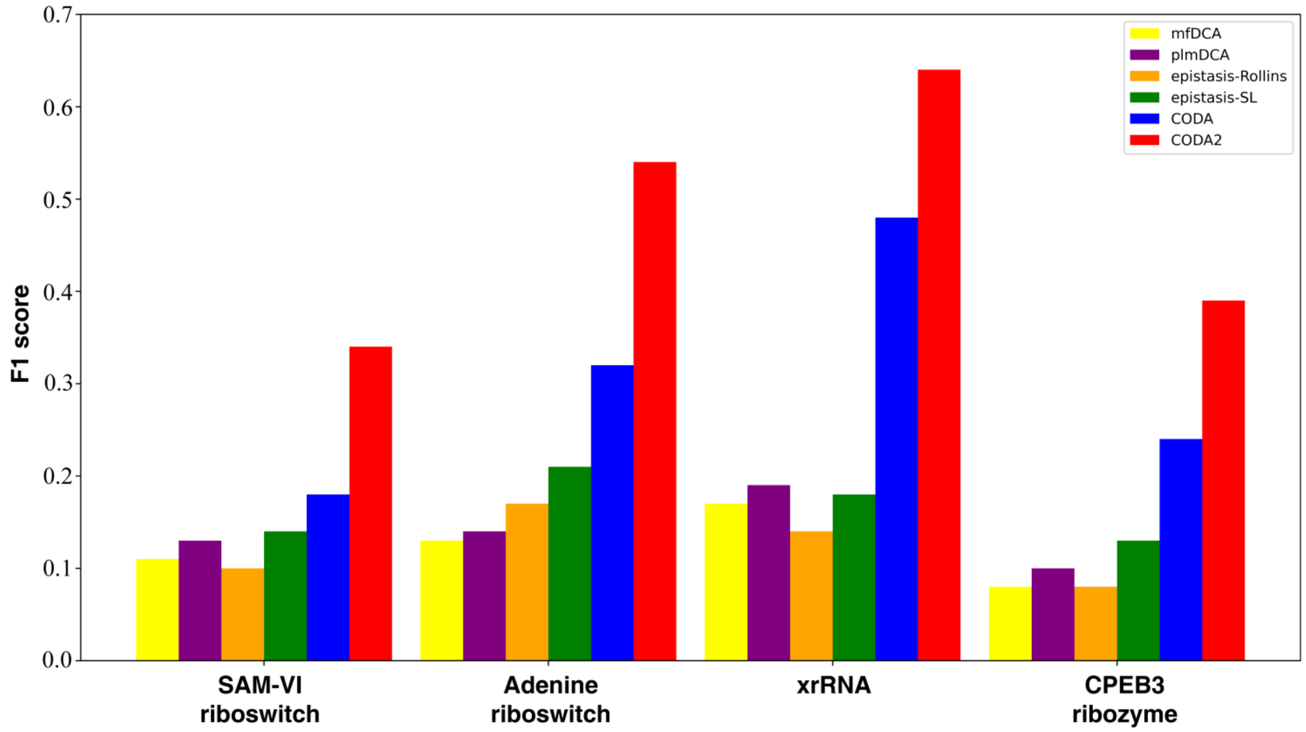


**Supplementary Figure S2. Improvement CODA2 over CODA and other related techniques.** Comparisons of *F1* scores inferred by the present (i.e., CODA2) and previous version of the CODA[1] as well as the mean-field direct coupling analysis (mfDCA-RNA)[3], pseudolikelihood maximization coupling analysis (plmDCA-RNA)[3], epistasis from Schmiedel & Lehner (epistasis-SL)[4] and from Rollins *et al* (epistasis-Rollins)[5] for the SAM-VI riboswitch, adenine riboswitch, xrRNA, and CPEB3 ribozyme.

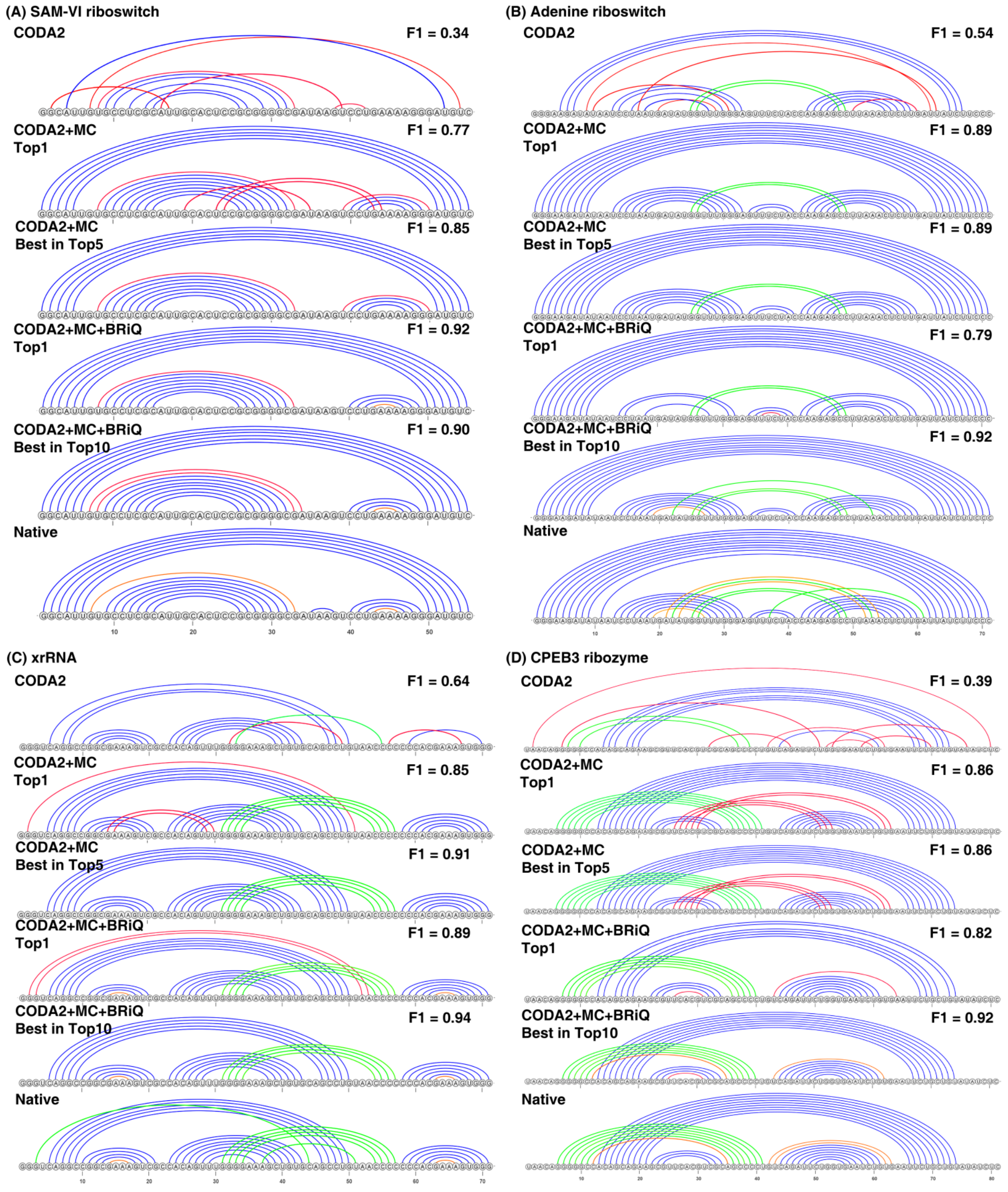


**Supplementary Figure S3**. **Capture of tertiary base pairs (pseudoknots) by MobiSeq**. The comparisons between base-pairs derived from CODA2 (top 1), CODA2+MC (top1), CODA2+MC (best in top 5), CODA2+MC+BRiQ (top1), CODA2+MC+BRiQ (best in top 10), and native structures for SAM-VI riboswitch (A), adenine riboswitch (B), xrRNA (C), and CPEB3 ribozyme (D). Native Watson-Crick base-pairs are shown in blue, and non-Watson-Crick base-pairs are shown in orange, except those pseudoknots (tertiary base pairs) indicated by green. False positive predictions are shown in red.

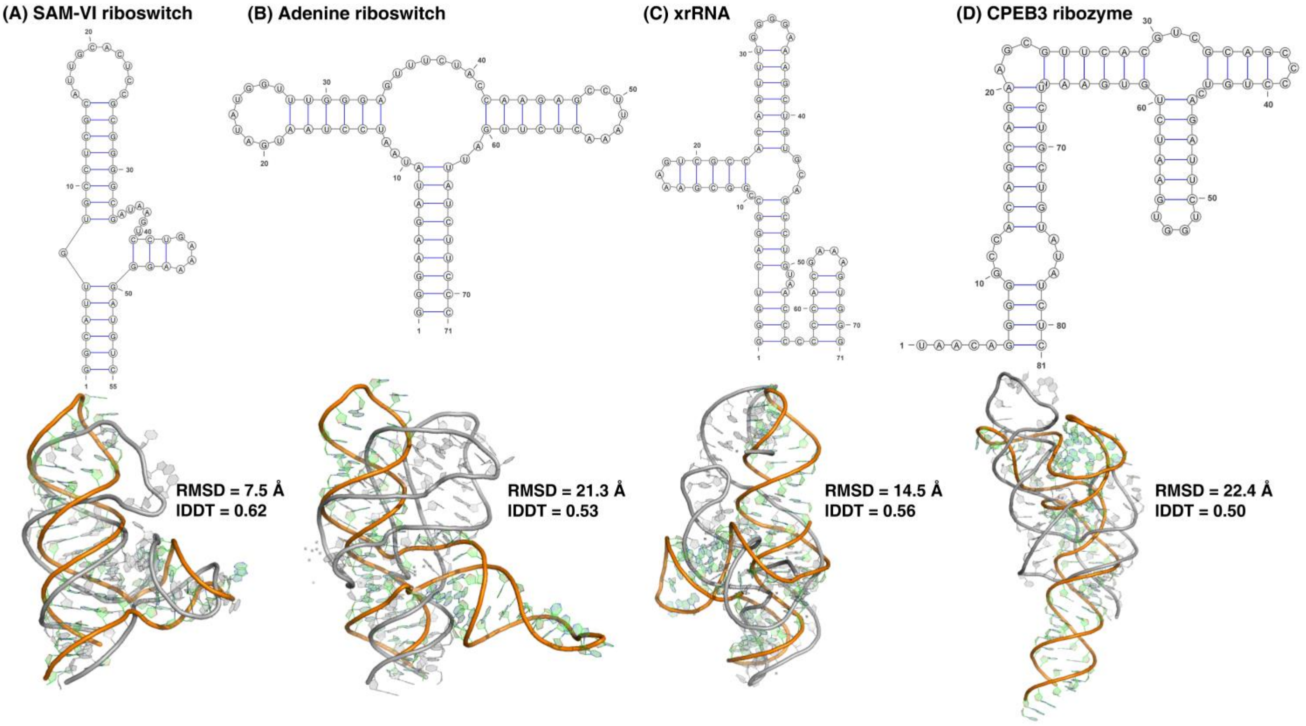


**Supplementary Figure S4**. **The results from RNAfold as the baselines.** Secondary structures (top) predicted by RNAfold[6] as well as the top 1 of 3D structures modeled by BRiQ using the predicted secondary-structures as restraints (bottom; grey: experimental structures; color: predicted structures.) for SAM-VI riboswitch (A), adenine riboswitch (B), xrRNA (C), and CPEB3 ribozyme (D).

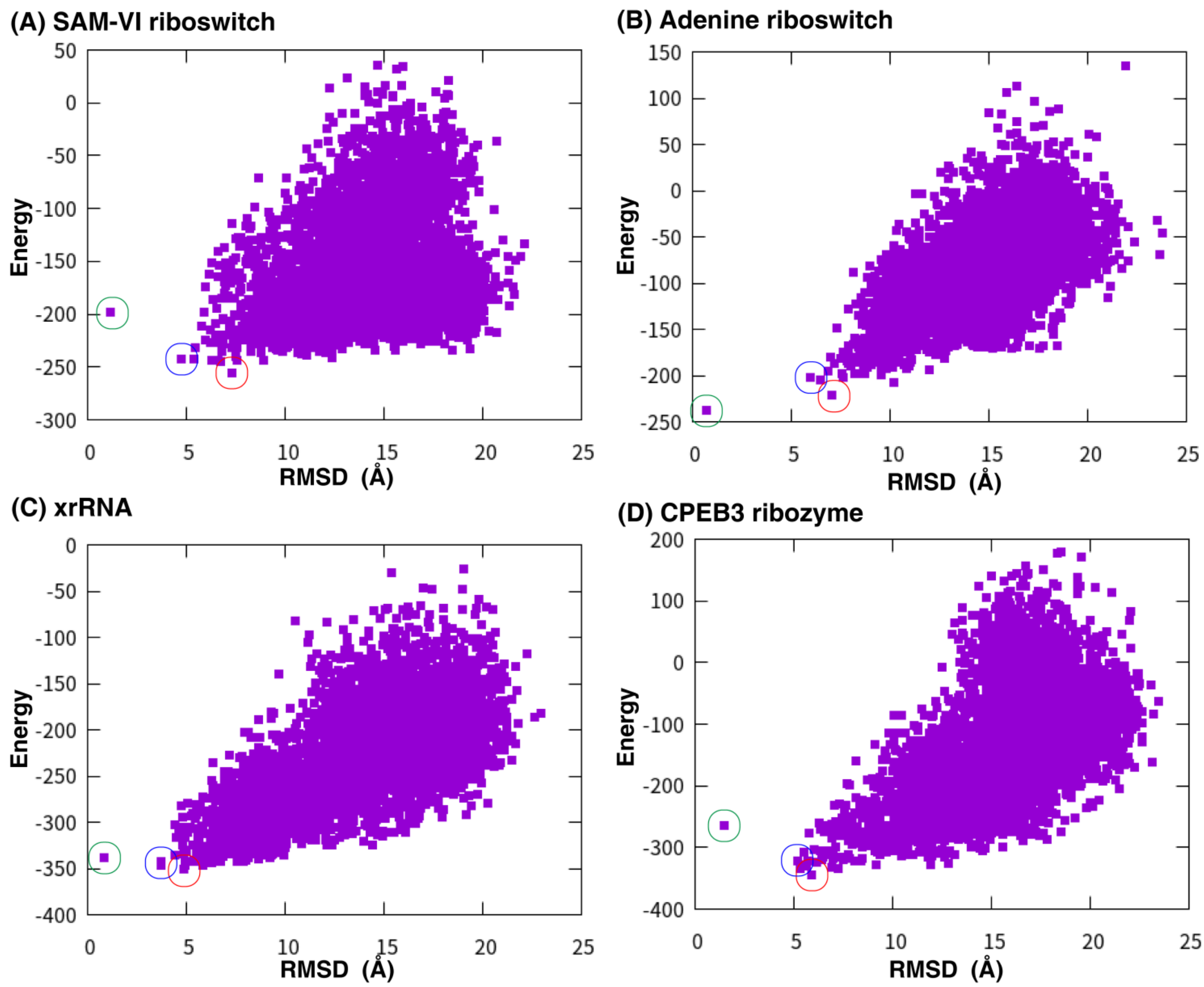


**Supplementary Figure S5**. **Predicted 3D models evaluated by BRiQ energy.** BRiQ Energy versus RMSD values of the conformations sampled by BRiQ[7]using the top 5 base-pairings inferred by CODA2+MC as the restraints for SAM-VI riboswitch (A), adenine riboswitch (B), xrRNA (C), and CPEB3 ribozyme (D). The points for top 1, best in top 10 and refined native structures are marked by red, blue and green circles, respectively.

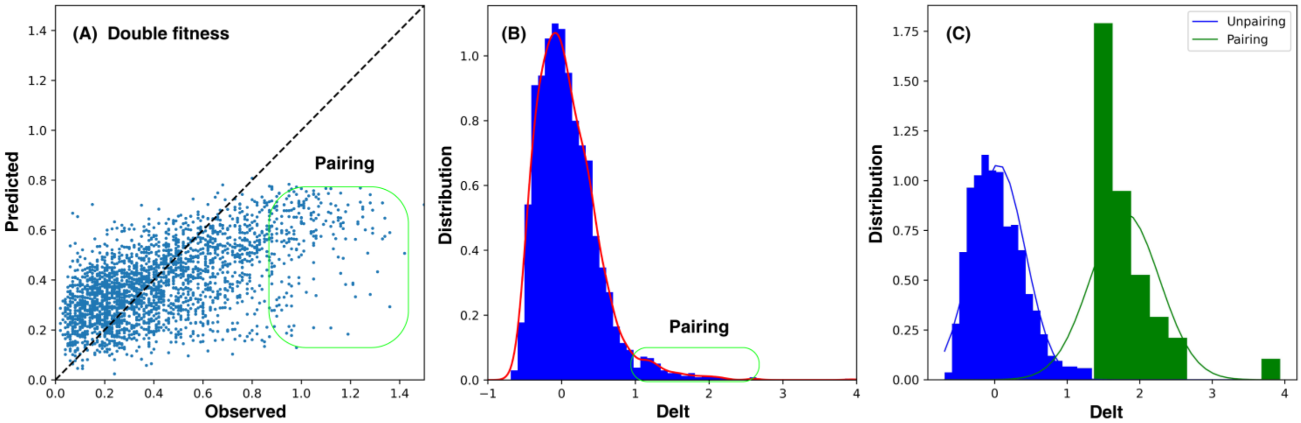


**Supplementary Figure S6. A schematic principle of classification in CODA2 .** (A) The fitness of double mutants observed from sequencing versus those predicted by the independent-mutation model for xrRNA. Outliers for likely base pairs (co-variated) are indicated by the green box. (B) The distribution (blue bar) as well as the kernel density estimation curve (red line) of the difference (i.e., Delt) between observed and predicted values of double fitness, and the distribution of possible base pairing is marked by the green box. (C) The data of Delt shown in (B) can be divided into two contributions representing whether they are likely to form pairs, according to whether the data is more than $n * \sigma$ (σ is the standard deviation of data, and *n* is 3.5 here), and the distributions of these two sets of data along with their corresponding normal distribution curves are shown here.
